# Evaluating Large Language Models as Tools to Navigate Researchers in Rapidly Evolving Research Landscapes: A Case Study in Cancer Drug Response Prediction

**DOI:** 10.64898/2026.09.02.748827

**Authors:** Elena Mourelatou, Ioannis Katakis

**Author notes:** This manuscript is the extended version of the work presented as a short paper at the 2025 IEEE/WIC International Conference on Web Intelligence and Intelligent Agent Technology (WI-IAT). At the time of this posting, the conference proceedings have not yet appeared in IEEE Xplore.

## Abstract

Large Language Models (LLMs) have emerged as promising tools for assisting researchers in automating and accelerating the synthesis of literature reviews. However, their reliability is a significant concern due to issues like factual inaccuracies and hallucinations. The key question is whether LLMs can reliably provide comprehensive, up-to-date overviews and analyses. This study evaluates the performance of three leading LLMs (OpenAI’s ChatGPT, Google’s Gemini, and DeepSeek) on the complex task of generating a comprehensive survey paper on deep learning for cancer Drug Response Prediction (DRP). By testing both standard and Deep Research (DR) / Deep Think (DT) modes of LLMs with prompts of varying detail, this paper assesses key academic dimensions, including reference management, content quality, and analytical depth. Key findings reveal that while DR modes of LLMs significantly improve reliability by eliminating hallucinations, performance variations exist across models and prompts. A trade-off between reference quantity and integration quality was observed, and even the best-performing models lacked the analytical depth of human experts, often requiring extensive human supervision. The study concludes that LLMs currently serve as powerful assistive tools but still cannot replace the critical validation and synthesis provided by human researchers. Choosing the best LLM to use depends on the task in hand, while several strategies can be implemented to improve the produced output.

## 1. Introduction

Research fields that are rapidly evolving, such as DRP in cancer therapeutics with Machine Learning and especially Deep Learning in the context of precision oncology, pose a challenge for researchers to stay up to date with the vast and exponentially increasing number of publications. In this field, various network architectures and learning strategies are being implemented using a variety of input data for drugs and cancers, with the goal of predicting cancer response to various treatments. These predictions can then be used to help clinicians in choosing the appropriate treatment for every individual patient. The complexity of the specific research field lies in the variety of the approaches implemented (in the architectures of learning schemes, data types – inputs and outputs-, and their combination), as well as the lack of a unified framework for performance evaluation and comparison between the different approaches. As a result, the manual task of identifying and analyzing the already tested approaches to find the most promising practices is challenging. Literature reviews, although valuable, are extremely time-consuming, and during the time required for the paper to be published (often more than a year) significant new research has already emerged, often rendering them outdated [1]. As a result, their utility for researchers who want to stay constantly updated is limited. Additionally, the process required for creating literature reviews involves a series of steps, which many times are affected by human bias and limitations [2], thus compromising their reliability. These issues underline the necessity for reliable automated tools that can assist researchers in the complex process of selecting, evaluating and synthesizing information from the exponentially expanding scientific literature.

The recent evolution of LLMs offers tremendous capabilities that can enhance various aspects of scientific research, including literature synthesis and scientific writing [3]. In an attempt to reduce their cognitive load, researchers are exploring LLMs as tools to automate the compilation of literature reviews [4], [5], [6]. This way, they could focus on strategically designing next steps, based on a comprehensive and current understanding of the work performed so far (including the latest findings), thereby accelerating scientific discovery.

Although this potential of LLMs sounds promising, their contribution lies in the reliability and trustworthiness of their outputs. However, there are several concerns raised by the scientific community related to LLMs’ responses, with the most pronounced ones being “hallucinations”, which are plausible but fabricated information [7], and generalization bias, in which LLMs tend to make conclusions broader than what is supported by the source [8]. These limitations can lead to inaccurate and misleading information, thus threatening scientific integrity.

This study provides an extensive evaluation of three commonly used LLMs (OpenAI’s ChatGPT, Google’s Gemini and DeepSeek) on their ability to generate comprehensive and scientifically accurate surveys in the rapidly evolving research field of DRP for cancer treatment. These models are tested in their default mode as well as their DR or DT mode, designed for more in-depth analysis and reasoning. Various prompt conditions are tested for writing the scientific survey paper with varying levels of guidance (minimum, medium, detailed, directed writing). For each prompt generated by each model, an analysis of how references are handled, as well as the quality of the content of the text generated by the LLMs is performed.

The **key research objectives** addressed in this study are:

1. Evaluating ChatGPT’s, Gemini’s and DeepSeek’s ability to generate a survey paper in a rapidly evolving research area such as DRP in cancer treatment.
2. Comparing the performance of the LLMs’ DR/DT modes against their standard versions.
3. Assessing the impact of varying prompt details on the responses generated
4. Assessing the LLMs’ ability to extract and synthesize information from uploaded documents for writing specific survey sections.
5. Identifying the best performing model in this task

### Contribution

This study provides a systematic and critical assessment of the practical utility of LLMs in generating a credible and academic useful survey focused on the complex field of DRP in cancer treatment, a field that has not been studied before in this context. Apart from the standard versions of the models, the newly developed DR / DT modes of the models, designed to offer more indepth analysis, are evaluated and compared with the standard modes, with very few studies in the literature including these modes. This is the first study to directly compare these three models, and their DR/DT modes, in such academic tasks. Additionally, this study provides a structured analysis of the strengths and weaknesses of each model tested, when prompts of varying levels of detailed instructions were provided and multiple prompts were used in more than one cases to provide feedback to the LLMs and improve the provided response. This approach differentiates the current study from other studies performed in this area. The overall findings aim to guide researchers on how to optimize the use of these tools, while being aware of their inherent limitations, and highlight how to combine LLM capabilities with human expertise to enhance productivity and accelerate scientific research.

## 2. Related Work

LLMs have been studied as tools to assist researchers in several steps of the literature review process, such as identifying research gaps [5], searching for relevant papers [6], [9], [10], assisting in the inclusion or exclusion process of publications by assessing their relevance based on their titles and abstracts [4], [9], [11], ranking retrieved papers based on their relevance with the abstract or keywords used [12], extracting data from the retrieved or provided papers [4], [10], draft sections of manuscripts (e.g., abstract, introduction, discussion, results) [4], as well as generating text by synthesizing information from multiple studies, with improved performance when they are guided to generate sections of a review by a specified plan or structure [6], [13]. The models’ performance in the various processes depends on the task as well as model architecture [9], [12].

Despite the promising capabilities of LLMs in academic writing, several limitations and challenges are presented, which threaten factual integrity, reproducibility, and analytical depth. The most significant issue is hallucinations, meaning the generation of plausible but non-factual information, especially in citations and references which are the cornerstone of academic integrity [14]. In the context of literature reviews, LLMs can either create context that is not included in the paper cited in text or create publications that do not exist (i.e., authors, DOIs, paper title etc. are not real) [12]. Hallucination rates can be significantly high, thus threatening the accuracy and reliability of scientific work [15] .

Another challenge faced when using LLMs for academic writing is their underperformance in terms of critical analysis and insightful synthesis, as well as identifying gaps for future research, especially when less structured approaches are used[5]. Moreover, their reproducibility poses significant challenges, since the same prompt can yield different outputs, even if it was used in the same model in subsequent runs, due to their probabilistic nature, as well as the continuous updates performed in commercial closed-source models which lead to changed behavior of LLMs over time. These issues make it impossible for other researchers to replicate the results produced in a study. Additional challenges include the bias that LLMs inherit from the vast amount of text used for their training [9], the fact that they are not always able to access the latest publications or papers published in peer-reviewed journals that have subscriptions, which can lead to outdated or incomplete knowledge [16], the restrictions in the LLMs’ processing of lengthy inputs, such as full scientific manuscripts[11], and the substantial demands on computational power and infrastructure required when a large number of lengthy documents needs to be evaluated [9].

To address the above-mentioned challenges and be able to fully exploit the potential of LLMs in academic writing, researchers have implemented specific methodologies and frameworks, such as Retrieval-Augmented Generation (RAG), where information inputted to LLMs are collected from an external knowledge database thus increasing reliability, refinement through feedback and reasoning [17], and prompt engineering, where providing prompts with specific structure and more details can refine the LLMs response. Specifically, in the context of literature review, better results are achieved when breaking down the process in smaller sub-tasks, instead of providing a single request and then reviewing and revising the outcome [5].

Several studies compare the performance of different LLMs in the task of academic writing, highlighting their strengths and limitations across different versions and specific tasks. Gemini (formerly known as Bard) was found to have superior performance compared to ChatGPT-3.5 in scientific writing, in terms of reference formatting and inclusion of detailed explanations from scientific papers [18]. Later versions, such as Gemini 2.5 Pro, had an improved performance in terms of hallucinations and literature search [19], however it produced more generalized outlines compared to ChatGPT in data extraction and abstract generation [18], [19]. In the ChatGPT family, o4-mini had a better performance in retrieving relevant papers without hallucinations, however it underperformed compared to human researchers and showed high plagiarism rates for paraphrased abstracts [19], [20], [21]. Moreover, ChatGPT provided more creative and elaborate title suggestions and detailed outlines [18]. On the other hand, DeepSeek V3 exhibited a content volume similar to ChatGPT, with moderate rates of plagiarism [22]. Its Deep Think R1 module outperformed Gemini 2.5 Pro Experimental as well as ChatGPT o4-mini-high in terms of correct entries in data extraction for systematic reviews, although minor server issues and requirement for complex procedures to upload papers were observed [19].

The **research gap** that our paper aspires to fill is comparing DR/DT and standard modes of commonly used LLMs in the task of survey paper generation, for ultimately evaluating their utility as research assistants in rapidly evolving research fields, using as a case study DRP in cancer treatment. This complex research area hasn’t been included in this type of studies before.

## 3. Methodology

Three different LLMs were evaluated in terms of their ability to generate a survey paper in the field of DRP in cancer treatment. Specifically, ChatGPT 4o (GPT-4), Gemini (2.0 Flash and 2.5 Flash) and DeepSeek-V3. In all LLMs, apart from the standard mode, the DR or DT (depending on the model) mode was also tested and evaluated. ChatGPT and Gemini were chosen since they are the market leaders and their recently released DR modes offer increased reliability and in-depth analysis in academic and research-oriented tasks. DeepSeek’s selection was based on the fact that this is a newer open-weight model, which has gained recognition due to its strong performance in coding and reasoning tasks. Each LLM has distinct technical architecture and varied strengths providing a foundation for an analysis on the capabilities that are more valuable in this type of academic work.

Five different prompts were used (see Table 3.1), to see the differences obtained in the LLMs’ responses depending on the context and level of detailed instructions provided by the user in the prompt. Three different levels of detail in the instructions provided in the user prompts were assessed.

**Table 3.1.** Prompts Used: A SERIES OF PROMPTS WITH INCREASING LEVELS OF DETAIL.

| **General Objective:** Generate a survey paper in the field of drug response prediction in cancer treatment. |  |  |
| --- | --- | --- |
| **Prompt** | **Level of detail** | **Prompt content** |
| **P1** | **Minimal** | Title only |
| **P2** | **Medium** | Abstract provided with some content requirements (analyze developed DL models, identify patterns and prevalent methods, use publications up to August 2022, include summary plots) |
| **P3** | **Detailed** | Title, specific section-by-section outline, and instructions (use only peer reviewed publications, no review articles - only original research) published from 2013 up to August 2022, Harvard referencing style, include tables and figures, use academic tone) |
| **P4** | **Focused – complex scenario** | Write “Dense Layers” section from uploaded PDFs as part of the survey (instructions: use academic style, in text citations in academic format, specific word count – around 250 words- requested in follow up prompt) |
| **P5** | **Focused – simple scenario** | Write “Graph Structures as drug representation” section from uploaded PDFs as part of the survey (instructions: use academic style, in text citations in academic format, specific word count – around 250 words) |

The responses obtained from each LLM were evaluated in terms of a) referencing handling and b) content quality (Table 3.2). The structure and content of the LLMs responses was compared to the one used in a reference review paper published in 2023 [23], that was considered to be more informative and helpful for researchers new to the specific field.

**Table 3.2.** Evaluation Criteria.

| Criteria for reference handling | Criteria for content quality |
| --- | --- |
| R1. Prompt dependency for reference inclusion<br>R2. Quantity<br>R3. Reliability:<br>● hallucinations<br>● irrelevant references<br>R4. Referencing accuracy<br>R5. Types of sources used:<br>● research papers<br>● review papers<br>● peer or non-peer reviewed publications<br>R6. Timeliness of sources:<br>● Usage of recent or older publications. Three timeframes were used:<br>○ < 2020<br>○ 2020 - 2022<br>○ > 2022<br>● Adherence to time frame provided in prompt | C1. Structure (logical structure – comparison with reference review)<br>C2. Style (writing quality, grammatical accuracy, flow)<br>C3. Length / analysis depth (coverage of key points from the reference review, providing enough analysis and examples of models, clarity)<br>C4. Correct information / referencing<br>C5. Summary plots / tables |

When Prompt 3 was used in ChatGPT 4o, two different versions were generated by the model (3vs1 and 3vs2), as part of the Reinforcement Learning from Human Feedback (RLHF) used in this model for fine-tuning. Both of these versions were kept and analyzed. Moreover, in Gemini, when the standard versions were used (2.0/2.5 Flash), a survey paper was not generated in Prompts 2 and 3. Instead they provided the structure of the survey paper and instructions on how to do it. This was attributed to the more detailed instructions included in these Prompts and the inherent limitations of the specific models to perform complex tasks, as well as an honesty in providing instructions and outline instead of a problematic, in terms of reliability, text.

The best performing versions of LLMs based on the initial evaluation performed on the responses of Prompts 1-3 were asked to generate specific sections of the survey paper based on a small number (8-10) of uploaded papers (**Prompts 4 and 5**). These prompts tested the ability of LLMs to extract, synthesize, and summarize information based on specific literature, instead of relying on general knowledge or web search, tasks that are required in academic reviews. Two different topics were chosen to test the LLMs performance in a complex and a simpler scenario. The first topic was the “Dense Layers” subsection in Neural Network modules for DRP of the reference review paper (Prompt 4). In this topic, which is considered more complex, an analysis of the specific methodology with several model examples is required. The second topic was “Graph Structures”, which is a subsection in the Representations of Drug Compounds section (Prompt 5). In this simpler scenario, an analysis of the advantages of this type of drug representations compared to other types is required. The evaluation of LLMs performance in Prompts 4 and 5 was based on limitations in the uploading procedure, adherence to word count constraint, quality of synthesized information and the completeness of model descriptions (where available), as well as usage of academic format for references.

The models were queried with Prompts 1-3 at the beginning of May 2025 in their respective web UI (i.e., chat.openai.com for ChatGPT, gemini.google.com for Gemini and chat.deepseek.com for DeepSeek). By the end of May, the Flash 2.0 model in Gemini was no longer available. As a result, Prompts 1-3 were used to query Flash 2.5 in Gemini at the end of May. Prompts 4 and 5 were used to query all models in the middle of June 2025.

## 4. Results AND Discussion

### 4.1 Referencing Handling Evaluation of LLMs’ responses to Prompts 1-3

The responses of the LLMs produced from each of the used prompts were initially evaluated in terms of reference handling (criteria R1-R6 in Table 3.2). Standard versions of all three LLMs generally did not include references or in-text citations unless explicitly prompted to do so (Table 4.1). In some cases, an additional prompt to include in-text citations (apart from reference list) was needed. In contrast, the DR modes of ChatGPT and Gemini autonomously included references and in-text citations, attributed to being designed and fine-tuned to perform in depth multi-step research for complex tasks. Thus, reference inclusion is necessary to increase the credibility of their responses and produce higher level research-oriented outputs. DeepSeek’s DT mode, however, still required a specific prompt to include references, since its primary purpose is reasoning for tasks requiring rigorous logic.

**Table 4.1.**
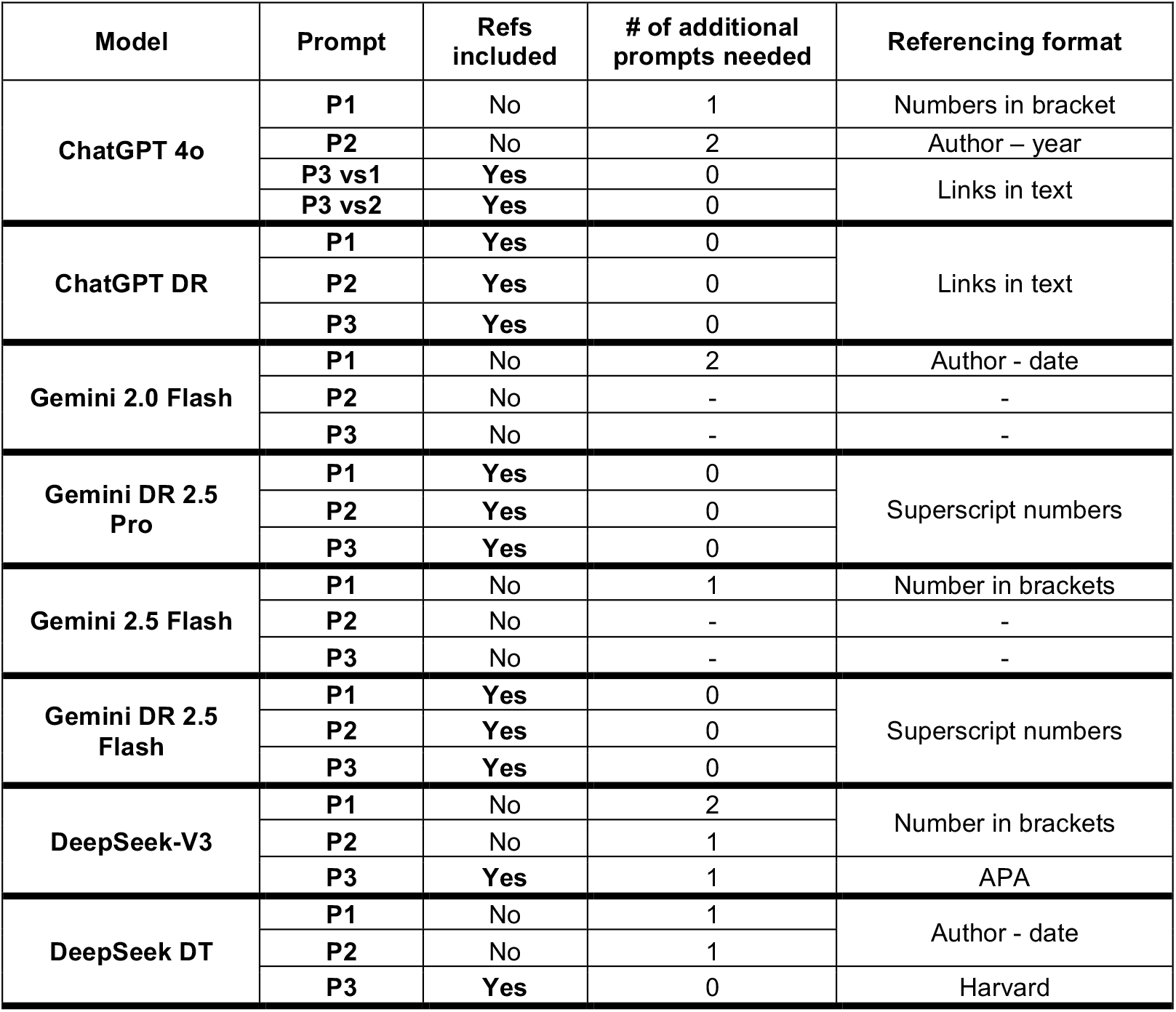
LLMS Ability TO INCLUDE References Autonomously AND Referencing Style.

| Model | Prompt | Refs included | # of additional prompts needed | Referencing format |
| --- | --- | --- | --- | --- |
| ChatGPT 4o | P1 | No | 1 | Numbers in bracket |
|  | P2 | No | 2 | Author – year |
|  | P3 vs1 | Yes | 0 | Links in text |
|  | P3 vs2 | Yes | 0 |  |
| ChatGPT DR | P1 | Yes | 0 | Links in text |
|  | P2 | Yes | 0 |  |
|  | P3 | Yes | 0 |  |
| Gemini 2.0 Flash | P1 | No | 2 | Author - date |
|  | P2 | No | - | - |
|  | P3 | No | - | - |
| Gemini DR 2.5 Pro | P1 | Yes | 0 | Superscript numbers |
|  | P2 | Yes | 0 |  |
|  | P3 | Yes | 0 |  |
| Gemini 2.5 Flash | P1 | No | 1 | Number in brackets |
|  | P2 | No | - | - |
|  | P3 | No | - | - |
| Gemini DR 2.5 Flash | P1 | Yes | 0 | Superscript numbers |
|  | P2 | Yes | 0 |  |
|  | P3 | Yes | 0 |  |
| DeepSeek-V3 | P1 | No | 2 | Number in brackets |
|  | P2 | No | 1 |  |
|  | P3 | Yes | 1 | APA |
| DeepSeek DT | P1 | No | 1 | Author - date |
|  | P2 | No | 1 |  |
|  | P3 | Yes | 0 | Harvard |

The **referencing styles used** in the LLMs responses varied across models (Table 4.1), and in some cases across prompts (e.g., ChatGPT 4o, DeepSeek). DeepSeek’s DT mode was the only one to use Harvard style in Prompt 3, which was the referencing style specifically requested in the prompt. This lack of consistency in referencing formatting and inability to use a requested referencing style poses a significant challenge for academic writing.

In terms of **reference quantity**, Gemini DR, especially 2.5 Pro model (37 - 92 references), provided the most extensive reference lists (Fig. 4.1a). However, several of the references used were duplicates (up to 30%), since different URLs pointing to various repositories (e.g., PubMed, ResearchGate, or arXiv) were considered as different references (Fig. 4.1b). This indicates a retrieval pipeline that fails to perform effective deduplication. Moreover, not all references listed were cited in text (20-60.9% in DR 2.5 Pro and 46-90% in DR 2.5 Flash were actually cited), which makes it necessary to manually verify that all references are properly integrated in text. This indicates that although these models have an effective retrieval system, they have a poor ability to integrate them in text. And although the content may be factually correct, it is practically difficult to verify it. The smallest number of references was used by ChatGPT.

**Fig. 4.1.**
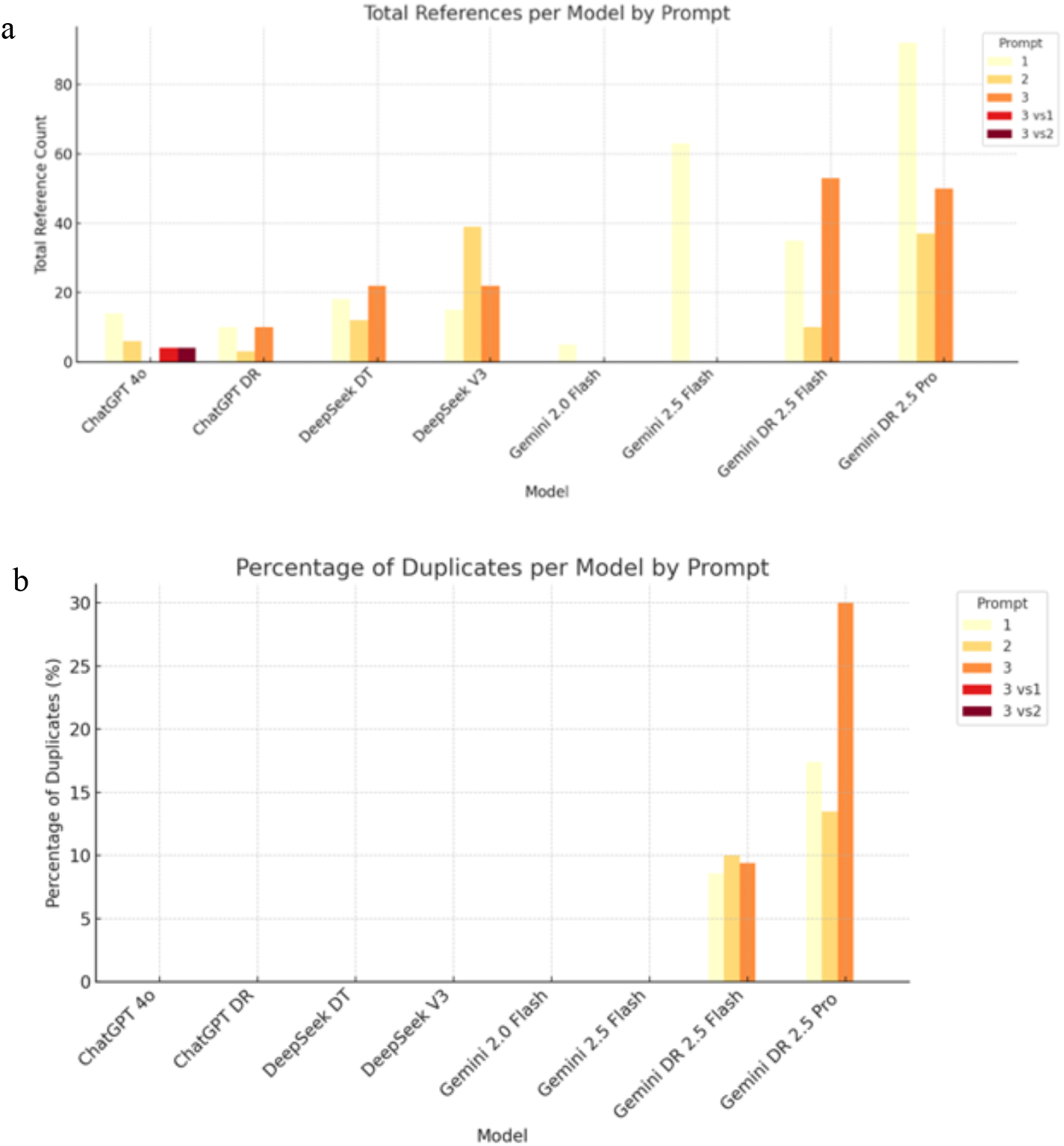
(a - top) Total references in each model by each prompt (b - bottom) Percentage of duplicate references produced by each model in each prompt.

However, quantity should be evaluated alongside with **reliability and accuracy** to have a more comprehensive analysis of each LLMs’ abilities. In this regard, the standard versions of Gemini and DeepSeek had high hallucination rates making them less reliable for academic tasks, while ChatGPT 4o produced hallucinations only in Prompt 2 (Fig. 4.2). This finding is aligned with relevant studies, where GPT-4 and Gemini were found to have a relatively high hallucination rate when prompted to conduct systematic reviews (28.6% and 91.4% respectively) [15]. Moreover, Gemini Advanced was found to produce higher hallucination rates in the provided references compared to ChatGPT-4o (76.7% versus 20.0%) when asked to generate financial literature reviews [24], a behavior also observed in this study with standard Flash models. The DR modes of ChatGPT and Gemini successfully produced zero hallucinations, a significant advantage for research-oriented tasks, whereas in DeepSeek’s DT hallucinations were reduced but not completely eliminated (Fig. 4.2). Generally, the high rates of fabricated references observed in DeepSeek, make its use for academic writing risky. These results are in accordance with the literature, where ChatGPT-4o and especially DR performed significantly better in terms of hallucinations in generating references compared to DeepSeek DT (39.14%, 26.57% and 91.43% respectively) [25]. However, in that study ChatGPT DR also produced hallucinations, although to a lower extent, a behavior that was not observed in the current study.

**Fig. 4.2.**
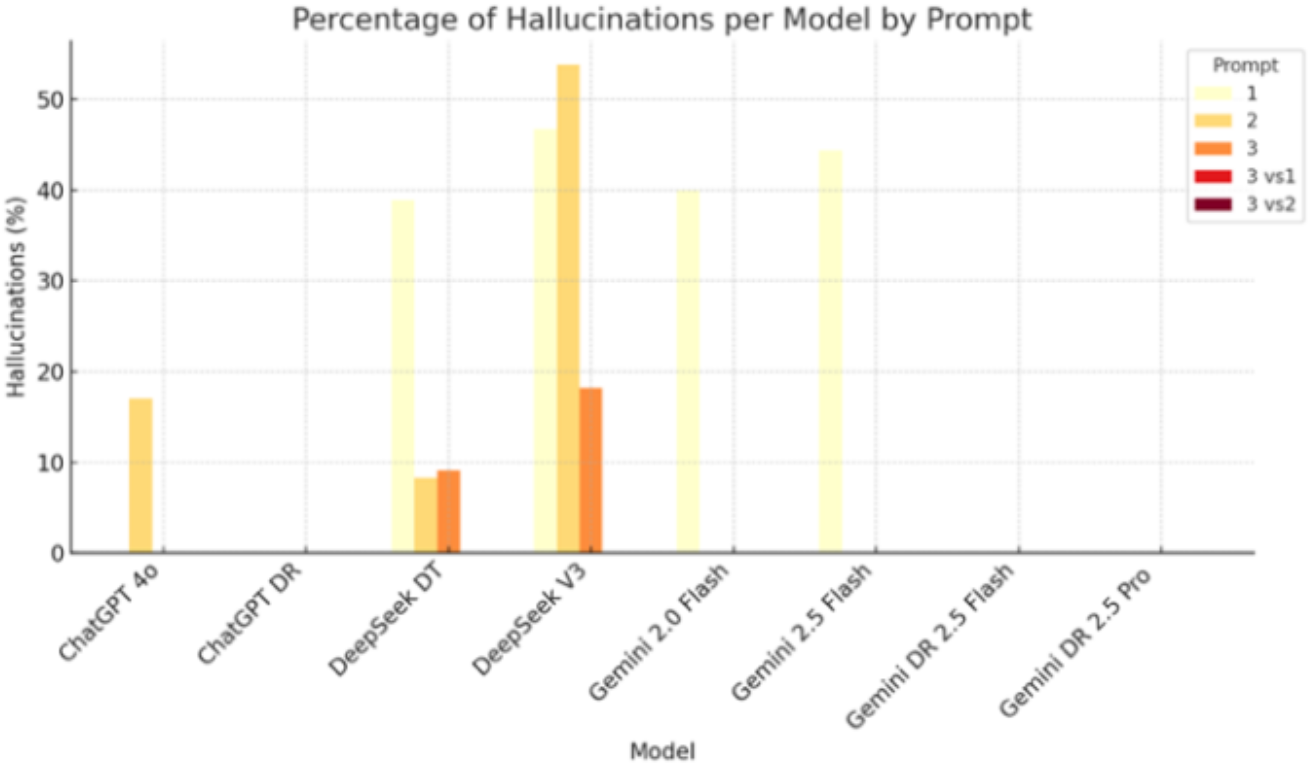
% of hallucinated references produced by each model in each prompt

Additionally, LLMs’ standard modes were more prone to include irrelevant or out-of-context references, with the exception of Gemini (Fig. 4.3). The highest reliability was demonstrated by ChatGPT DR, although it had the smallest number of references as mentioned above. Another disadvantage exhibited by DeepSeek in both modes was the poor referencing accuracy, exhibited with many mistakes in citation details (5.6 – 81.8%).

**Fig. 4.3.**
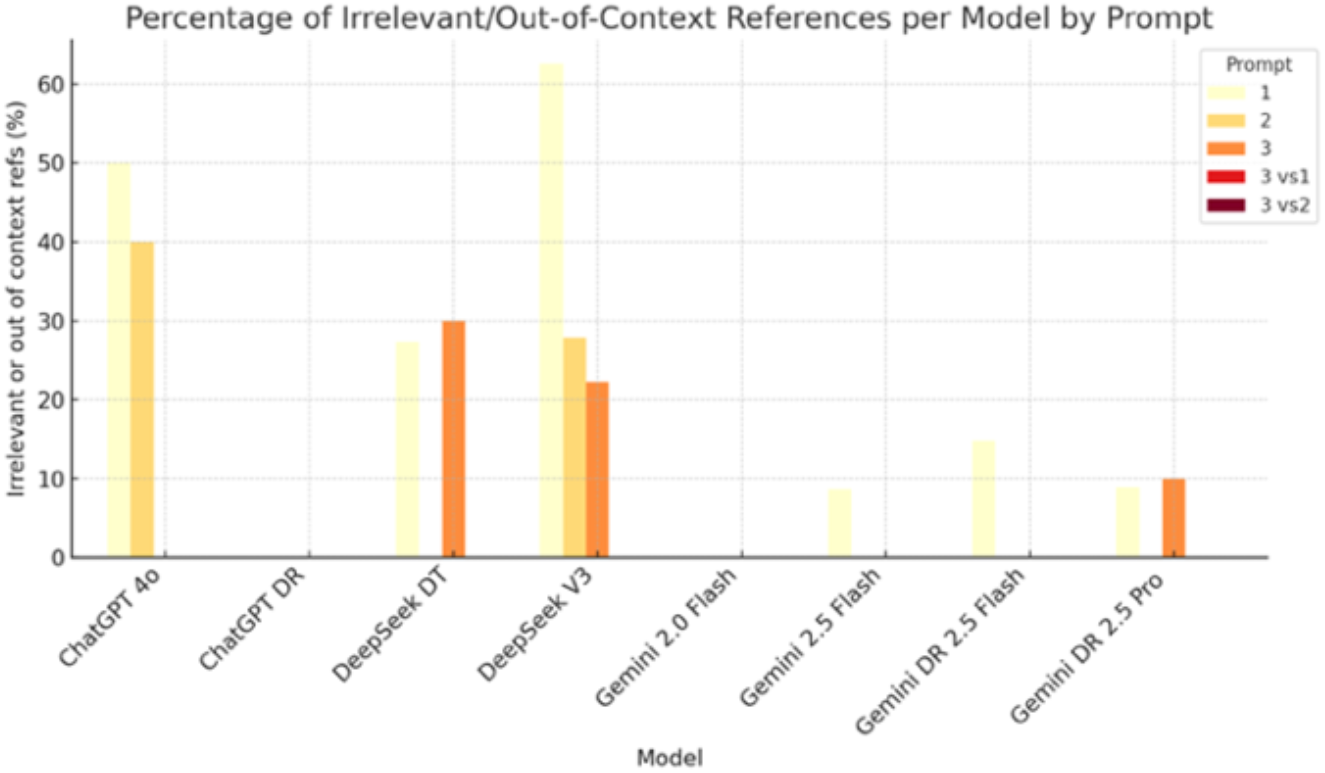
% of irrelevant references used by each model in each prompt

Most of the sources used by the LLMs in their responses were peer-reviewed papers, which are appropriate for a survey paper. Non-peer reviewed references included publications in arXiv and biorXiv (with several being published at a later stage), as well as various websites. Additionally, most of the papers retrieved and used by the LLMs in each prompt were research papers, a characteristic necessary for a survey paper intended to serve as a guide to a rapidly evolving research area. However, it should be noted that in some cases (e.g., ChatGPT DR P2 and Gemini 2.0 Flash P1 responses), the LLMs identified the provided reference review as a primary source of information and used this reference almost exclusively, with only a small number of research papers being additionally used. Furthermore, in Prompt 3 LLMs were asked to include only peer reviewed publications and original research papers (no review articles), a constraint that was only fulfilled by ChatGPT DR and DeepSeek-V3 in terms of peer reviewed publications, and DeepSeek DT and ChatGPT 4o in terms of research papers. This indicates DeepSeek’s advantage in being more successful in fulfilling prompt constraints, while other were unable to filter the source type.

In terms **of publications dates** of the used sources, Gemini and ChatGPT in DR modes showed a preference for recent publications, including papers from the current year (2025), while DeepSeek and Gemini 2.5 Flash tended to use older sources (Fig. 4.4), although their knowledge cut off was much later.

**Fig. 4.4.**
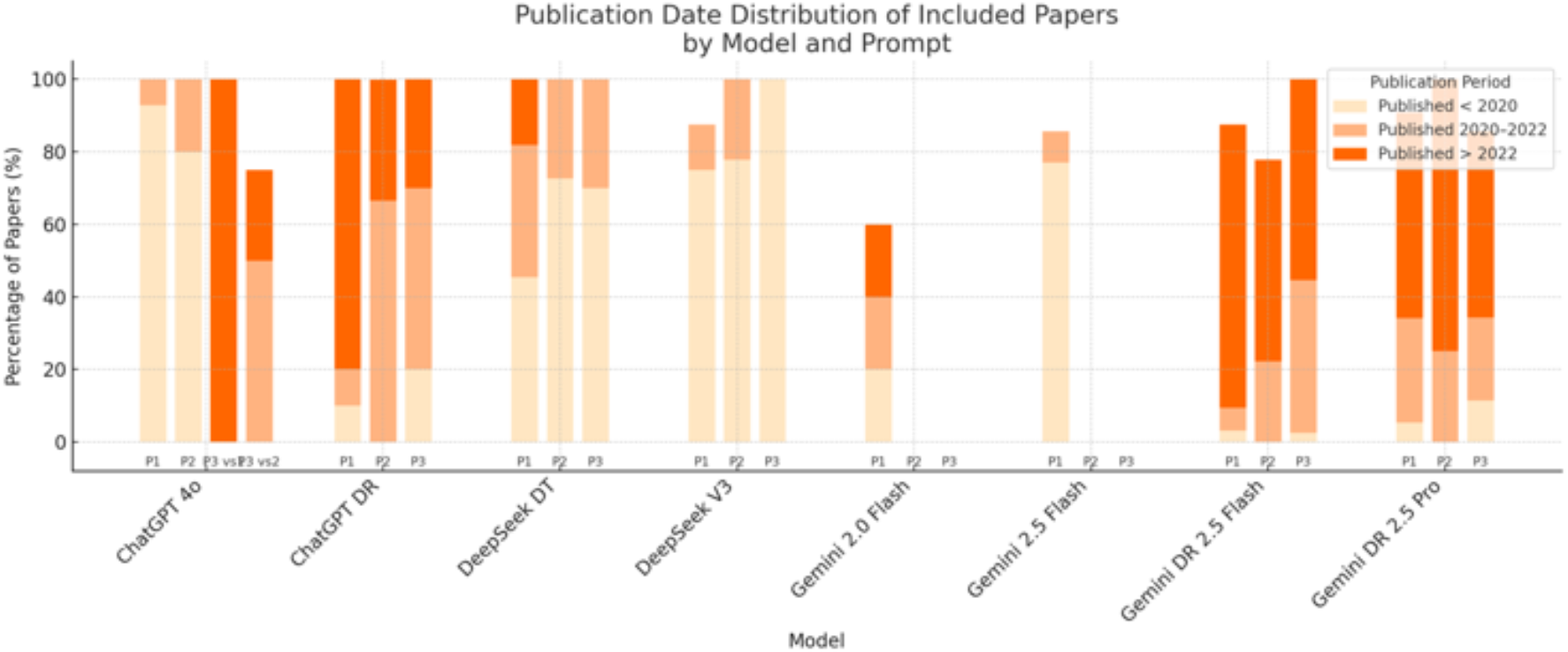
Publication date distribution of *papers* used as sources by each model in each prompt.

Time constraints were provided in Prompts 2 and 3, where it was requested to use papers published up to August 2022 for two reasons; (1) to be able to directly compare results and conclusions with those obtained in the reference review, having the same pool of publications, and (2) to exclude the reference review from the possibly retrieved papers hence obstructing the LLMs’ being influenced by it. However, several LLMs (ChatGPT 4o, ChatGPT DR, Gemini DR) included papers outside the specified timeframe. This indicates that while they can interpret the instructions, their retrieval mechanisms might prioritize relevance or internal knowledge structures over strict date adherence, or that they are unable to perform precise temporal filtering of the retrieved sources. However, although for the generation of survey papers focusing on contemporary advancements in a specific research field can be considered advantageous, in this study this behavior created a specific problem. Since the LLMs didn’t follow the publication date constraints provided in the relevant prompt, the reference paper was frequently included in the sources used by the LLMs, and due to its content matching the requested information, many LLMs (e.g., ChatGPT DR, Gemini DR 2.5 Pro and Flash in Prompts 2 and 3) relied heavily on this paper for content generation thus limiting the LLMs’ ability to conduct truly independent and comprehensive literature surveys. This indicates that they possibly prefer to re-process and synthetize any document that highly matches the provided description, thus simplifying the task in hand. As a result, the generated outputs lacked originality, often resulting in an overview of the information provided in the reference paper, rather than locate and synthesizing information from a broad range of relevant sources.

### 4.2 Content Quality Evaluation of LLMs’ responses to Prompts 1-3

The second part of the LLMs responses evaluation consisted of an analysis of the quality of the generated content to assess its academic utility (criteria C1-C5 in Table 3.2). For structural completeness, the structure of surveys generated by LLMs was compared to the one provided in the reference review. Generally, the level of detail in the prompt strongly influenced the structural completeness of the output. All of the models kept the detailed structure provided in Prompt 3. However, when less specific Prompts were used (i.e. P1), key sections found in the reference review were omitted in the generated surveys. Gemini 2.5 Flash outperformed 2.0 Flash, while DR modes of both ChatGPT and Gemini produced more comprehensive structures, while still missing some sections. Furthermore, Gemini in DR mode (in both 2.5 Flash and Pro) included topics not found in the reference review. This could be attributed to the fact that these models used mainly papers published after 2022, so they were exposed to probably other emerging trends than those identified in the reference review, which contained papers published up to August 2022. DeepSeek’s responses, in both the standard and DT modes, consistently omitted several key sections from their generated outlines, though they occasionally added relevant emerging topics. The observation that the model’s performance is highly sensitive to prompt design, and that using more detailed structured prompts is essential in generating high-quality responses, is in accordance with the relevant literature [26], [27].

In terms of **adherence to academic writing style**, most models, especially the standard versions as compared to the DR ones, employed frequently bullet point formatting in their texts, rather than continuous text, with little to no analysis provided in each section. Although bullet points are useful for summarizing or listing, their usage in academic writing, especially survey papers, is not that common and its extensive use is indicative of lack of analytical depth and ability to combine information to create a comprehensive text. This demonstrates that the standard versions of LLMs are mainly listing methods or concepts without being able to incorporate them into a critical academic discussion. Gemini DR 2.5

Pro was the most successful at generating text without bullet points, creating a more professional academic style. In ChatGPT 4o, several sections didn’t include relevant references, while in DR mode several sections were in bullet point formatting, a behavior that was also observed in Gemini DR 2.5 Flash. All DeepSeek’s outputs were considered brief overview reports rather than comprehensive academic surveys, where all information were delivered in bullet points with minimum or no analysis.

In terms of **depth of analysis**, the standard versions were found to systematically produced shorter texts that lacked the analytical depth required for a comprehensive literature survey. While ChatGPT and DeepSeek models in their standard versions generated texts below 800 words, the responses of Gemini in its standard versions (2.0/2.5 Flash) were more verbose, with responses around 1100 words and even 2762 in one case (Flash 2.5 P1). On the other hand, DR modes generated more extensive and sophisticated content with word counts ranging from 2,534 (ChatGPT DR, P1) to 7,706 (Gemini DR 2.5 Flash, P1). This behavior was not observed for DeepSeek’s DT, which maintained the same style as the V3, with small length and minimum analysis provided. However, even the best-performing models often failed to provide the required analytical depth, listing concepts without critical discussion or providing sufficient examples of research models.

In terms of the LLMs ability to **summarize findings in plots and / or tables**, it was found that all LLMs were not able to provide visualizations and mainly generated legends or descriptions of figures and plots when visualizations were requested in the prompt (P2 & P3). In some cases, figures were copied by the reference review (e.g., ChatGPT DR P2). Few models, and specifically Gemini DR models, were actually able to generate informative tables summarizing results across different prompts. This suggests that most LLMs are able to understand the concepts of data visualization and summarization of information in tables, but only few of them can generate actual tables, and only one can do this autonomously (Gemini DR 2.5 Pro). In general, they lacked the ability to generate novel, insightful plots or populate tables with synthesized data that goes beyond direct extraction.

Another important observation is that to achieve a high-quality academic output, using only one prompt is not sufficient. The necessity of using multiple follow-up prompts (iterative prompting), as in the experiments performed in this study, to adjust and improve the generated response is evident. When researchers provide feedback to the LLM or request corrections (e.g., correct errors, add missing information or elements, change of style), the outcome can be refined up to an extent. This process is similar to an automated process called self-refinement, where the LLM iteratively generates feedback on its own output and refines it to improve the final outcome [28].

### 4.3 Evaluation of LLMs’ responses to Prompts 4 & 5

In this task, the best performing versions / modes of LLMs based on the previous evaluations were used, i.e. DR / DT modes of all LLMs. Different limitations in file uploading were exhibited in each model. In Gemini Pro plan only 10 files at a time could be uploaded. In ChatGPT, only 3 files could be uploaded in basic plan, while in Plus plan a daily uploading of 10-15 files (50-100 MB per file) was permitted. DeepSeek has a reported file upload limit of 20 files per conversation with 512MB per file. However, when 8 pdf files were uploaded it stated that it could only read 56% of the uploaded files and could not proceed with the prompt. This was attributed to the dense text of research papers and the LLMs’ ability to process only a certain amount of text in each conversation. As a result, the processing limit of DeepSeek was exceeded. In order to overcome this issue, the papers were processed in batches of 2 or 3 at a time. DeepSeek was asked to analyze these fewer papers and generate an answer. After all the papers were processed this way, the generated texts from each batch were given back to DeepSeek to produce the final output.

In prompt 4, the LLMs initially failed to generate a text that had the typical length of a subsection in a survey paper. Instead, they produced lengthy texts (Table 4.2), with the structure of a survey paper. When specific instructions were given in regards to the text length in an additional prompt, they exhibited the ability to synthesize key information from the provided documents, but underperformed in terms of controlling the output length while at the same time maintaining comprehensive detail. The texts generated were lengthier in all cases, which indicates that the models prioritize generating more text, but without necessarily adding all of the critical technical information. The problem of controlling output length to satisfy prompt constraints, is a documented limitation of LLMs with specialized frameworks being developed and tested to resolve this issue [29], [30], [31].

**Table 4.2.** Word COUNT OF LLM RESPONSES TO PROMPTS 4 & 5.

| Model | Word Count |  |  |
| --- | --- | --- | --- |
|  | Prompt 4 |  | Prompt 5 |
|  | Initial re-sponses | After word count constraint |  |
| ChatGPT DR | 2289 | 322 | 249 |
| Gemini DR 2.5 Flash | 8094 | 298 | 559 |
| Gemini DR 2.5 Pro | 8431 | 395 | 199 |
| DeepSeek DT | 609 | 444 | 344 |

Higher level of information synthesis was observed in Gemini DR 2.5 Pro and DeepSeek DT, where model examples are incorporated into the text as part of a discussion regarding dense layers capabilities, leading to improved text flow and coherence. However, despite this strength, they all consistently failed to include all of the key information that would fully describe the models. This indicates that although the models have the ability to group related information based on common factors or architectures, which is valuable in establishing a narrative flow in a scientific text, they cannot provide technical details in a high degree of precision for each example or model. Moreover, in-text citations and a reference list were not included in all cases. Only DeepSeek DT used academic referencing, which is in line with the findings from Prompts 1-3.

In the simpler scenario of Prompt 5, two of the models, i.e. ChatGPT DR and Gemini DR 2.5 Pro, managed to satisfy word constraint, while the other two (DeepSeek DT and Gemini DR 2.5 Flash) produced longer texts (Table 4.2), indicating that their behavior depends on the topic. For this less complex topic, Gemini DR 2.5 Pro produced a condensed summary, while 2.5 Flash became wordier, requiring several additional prompts to condense its output. This indicates that 2.5 Pro performs better in synthesizing complex information. Regarding content, all models showed the ability to synthesize information from the provided papers, discussing the benefits of graph representations and comparing different approaches, however they failed to include complete technical details. In this regard, different prompts could be used, requesting specific content or technical details, to see if the LLMs’ performance could be improved.

### 4.4 Summary of Results and Guidelines for Researchers

Based on the evaluation performed, several differences in the reliability and content quality were observed not only between different models, but also between different versions and operation modes of models in the same family, as well as different prompts used. Generally, the DR/DT modes performed better than the standard versions of LLMs. Amongst those, ChatGTP and Gemini (2.5 Flash and 2.5 Pro) performed better than DeepSeek in the generation of the whole survey paper. Between these two, there is no single best tool, due to the fact that each model outperformed the other at specific parts but underperformed in other parts, making it important for researchers to choose the appropriate LLM based on their priorities. Specifically, in terms of reference reliability, the best performing model was ChatGPT DR (zero hallucinations and irrelevant references, all sources were peer-reviewed) but the quantity of those references was low. On the other hand, Gemini models in DR mode produced a higher number of references with zero hallucinations but some of them were irrelevant, while not all of them were cited in text. Additionally, in terms of timeliness of sources both Gemini and ChatGPT were able to provide recent publications. In terms of content depth and analysis, as well as structural coherence, ChatGPT DR and Gemini DR are comparable, with Gemini models being slightly better. All model responses provide most of the key elements required but frequently lack the required depth of analysis. However, Gemini provided lengthier texts with summarization tables and had a better text flow compared to ChatGPT. Finally, in terms of adherence to prompt constraints and academic referencing style, DeepSeek had the best performance, although underperformed in all the other categories.

However, when specific papers were provided for the generation of a subsection of the survey, DeepSeek DT performed better, especially in synthesizing information from different models as well as correct referencing formatting. An interesting observation in this task was that the models’ performance changed depending on the complexity of the subject provided. For example, Gemini DR 2.5 Pro performed better on the subsection of dense layers, where an analysis of different models and a synthesis of their architectures and characteristics had to be performed, while underperformed in less complex tasks, such as the summarization of information regarding graph structures as drug representations.

A proposed workflow is one that combines human expertise with the speed of LLMs (Fig. 4.5). The LLMs can perform the more laborious tasks of information processing and text generation, while humans can provide directions, iterative feedback and critical judgment to improve the final result. If the LLMs generate an initial draft of a literature review, then researchers should verify the citations and refine the critical analysis. Otherwise, to increase factual accuracy, specific sources/publications can be provided to LLMs by the researchers. This way, a combination of the strengths of researchers (i.e., domain expertise and critical thinking) and LLMs (i.e., speed of processing information and generating text) can produce papers superior to those produced only by LLMs.

**Fig. 4.5.**
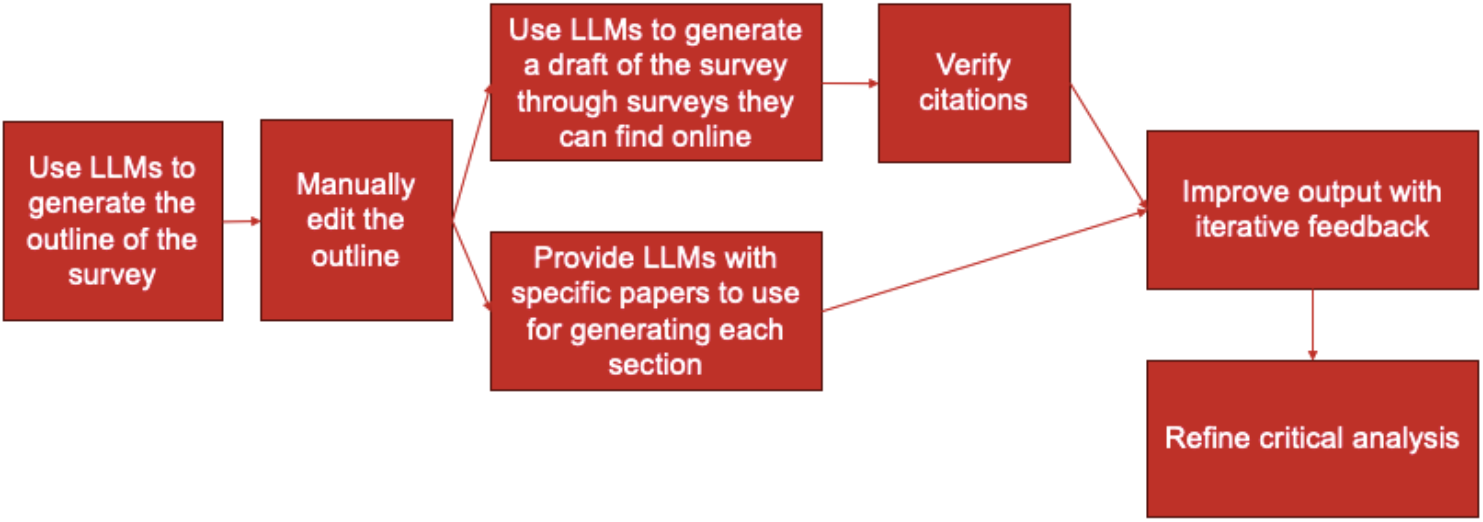
Proposed workflow for survey generation

## 5. Conclusions, Limitations, AND Future Work

Based on the evaluation performed in this study, the LLMs tested can act as powerful assistants but can not autonomously produce a high-quality, comprehensive, and fully accurate scientific survey in the field of DRP for cancer treatment. While they can significantly accelerate the drafting process, they cannot replace human critical thinking, validation, and synthesis. Choosing the best LLM to use depends on the task in hand, with DR modes found to be superior in terms of reliability and text generation. To imrpove the produced outcome several strategies can be implemented such as breaking the survey into subsections, providing the relevant literature for LLMs to use, providing specific structure and detailed instructions, as well as iterative feedback and corrections to guide the LLM, thus reducing the post-generation editing required. However, researchers should keep in mind that even when the above are implemented, models can still struggle with prompt constraints, and they should validate all information, verify every reference, and refine the content to ensure academic integrity and technical accuracy.

This study has three main limitations. First, it is focused on a highly specialized research area and the current findings may not be generalizable in other scientific disciplines. Second, the LLM landscape is rapidly evolving, with their performance changing at a very rapid pace. Hence, the performance of the specific models and versions evaluated in this study are snapshots in time. They may be quickly superseded by more advanced versions with different performance. Third, the content quality evaluation performed here was based on a comparison of LLM generated surveys with a single human-generated review paper, which involved a degree of personal judgement.

Future work could focus on enhancing LLM outcomes by developing frameworks that improve accuracy and adherence to academic standards. This includes further testing of advanced prompt engineering techniques and human-in-the-loop systems, which would integrate human expertise through dynamic feedback and refinement loops. Moreover, employing automated tools to evaluate LLM performance in survey generation would be valuable for accelerating the process and removing bias. However, the most significant challenge in automated evaluation would be assessing the novelty and analytical depth of the LLM-generated responses.

## References

[1] M. Sampson, K. G. Shojania, C. Garritty, T. Horsley, M. Ocampo, and D. Moher, “Systematic reviews can be produced and published faster,” J Clin Epidemiol, vol. 61, no. 6, pp. 531–536, Jun. 2008, doi: 10.1016/J.JCLINEPI.2008.02.004.

[2] N. R. Haddaway et al., “Eight problems with literature reviews and how to fix them,” Nature Ecology & Evolution 2020 4:12, vol. 4, no. 12, pp. 1582–1589, Oct. 2020, doi: 10.1038/s41559-020-01295-x.

[3] Z. Luo, Z. Yang, Z. Xu, W. Yang, and X. Du, “LLM4SR: A Survey on Large Language Models for Scientific Research,” ACM Comput Surv, vol. 1, p. 1, 2025, doi: 10.1145/nnnnnnn.nnnnnnn.

[4] D. Scherbakov et al., “The emergence of large language models as tools in literature reviews: a large language model-assisted systematic review,” Journal of the American Medical Informatics Association, vol. 32, no. 6, pp. 1071–1086, May 2025, doi: 10.1093/JAMIA/OCAF063.

[5] M. Zhao, F. Li, F. Cai, H. Chen, and Z. Li, “Can we trust LLMs to help us? An examination of the potential use of GPT-4 in generating quality literature reviews,” Nankai Business Review International, vol. 16, no. 1, pp. 128–142, Jan. 2024, doi: 10.1108/NBRI-12-2023-0115/FULL/XML.

[6] S. Agarwal et al., “LitLLM: A Toolkit for Scientific Literature Review,” Feb. 2024, Accessed: Jun. 28, 2025. [Online]. Available: https://arxiv.org/abs/2402.01788v2

[7] HuangLei et al., “A Survey on Hallucination in Large Language Models: Principles, Taxonomy, Challenges, and Open Questions,” ACM Trans Inf Syst, vol. 43, no. 2, Jan. 2025, doi: 10.1145/3703155.

[8] U. Peters and B. Chin-Yee, “Generalization bias in large language model summarization of scientific research,” R Soc Open Sci, vol. 12, no. 4, Apr. 2025, doi: 10.1098/RSOS.241776.

[9] F. Dennstädt, J. Zink, P. M. Putora, J. Hastings, and N. Cihoric, “Title and abstract screening for literature reviews using large language models: an exploratory study in the biomedical domain,” Syst Rev, vol. 13, no. 1, pp. 1–14, Dec. 2024, doi: 10.1186/S13643-024-02575-4/FIGURES/5.

[10] R. Peinl, J. B. Armin, H. Sarang, R. Chouguley, and S. Thalmann, “Using LLMs to Improve Reproducibility of Literature Reviews,” Association for Information Systems, Proceedings of the 2024 Pre-ICIS SIGDSA Symposium on Emerging AI Platforms for Societal Good, Dec. 2024, Accessed: Jul. 05, 2025. [Online]. Available: https://aisel.aisnet.org/sigdsa2024

[11] F. M. Delgado-Chaves et al., “Transforming literature screening: The emerging role of large language models in systematic reviews,” Proc Natl Acad Sci U S A, vol. 122, no. 2, p. e2411962122, Jan. 2025, doi: 10.1073/PNAS.2411962122/SUPPL_FILE/PNAS.2411962122.SAPP.PDF.

[12] S. Agarwal et al., “LitLLMs, LLMs for Literature Review: Are we there yet?,” Transactions on Machine Learning Research, vol. 2024-December, Dec. 2024, Accessed: Jul. 05, 2025. [Online]. Available: https://arxiv.org/abs/2412.15249v2

[13] C. C. Hsu et al., “CHIME: LLM-Assisted Hierarchical Organization of Scientific Studies for Literature Review Support,” Proceedings of the Annual Meeting of the Association for Computational Linguistics, pp. 118–132, 2024, doi: 10.18653/V1/2024.FINDINGS-ACL.8.

[14] D. Singh and A. Singh, “Ubiquity of LLM Hallucinations Across Critical Domains: A Survey,” pp. 115–132, 2025, doi: 10.1007/978-981-96-8197-6_9.

[15] M. Chelli et al., “Hallucination Rates and Reference Accuracy of ChatGPT and Bard for Systematic Reviews: Comparative Analysis,” J Med Internet Res, vol. 26, p. e53164. 2024, doi: 10.2196/53164.

[16] H. Letiche and M. Lissack, “Literature Reviews with Llm-Based Tools,” 2025, doi: 10.2139/SSRN.5110658.

[17] S. T. I. Towhidul et al., “A Comprehensive Survey of Hallucination Mitigation Techniques in Large Language Models,” ArXiv, Jan. 2024.

[18] H. S. AlSagri, F. Farhat, S. S. Sohail, and A. K. J. Saudagar, “ChatGPT or Gemini: Who Makes the Better Scientific Writing Assistant?,” J Acad Ethics, pp. 1–15, Jul. 2024, doi: 10.1007/S10805-024-09549-0/FIGURES/3.

[19] M. Sollini et al., “Human Researchers are Superior to Large Language Models in Writing a Systematic Review in a Comparative Multitask Assessment,” Jun. 2025, doi: 10.21203/RS.3.RS-6863946/V1.

[20] Ö. Aydin, E. Karaarslan, and F. Safa Erenay, “Generative AI in Academic Writing: A Comparison of DeepSeek, Qwen, ChatGPT, Gemini, Llama, Mistral, and Gemma,” 2025.

[21] I. Ul Haq Akhoon, M. Y. Khan, and T. A. Bhat, “Comparative Analysis of AI-Generated Research Content: Evaluating ChatGPT and Google Gemini,” Oct. 2024, doi: 10.21203/RS.3.RS-5265799/V1.

[22] Ö. Aydin, E. Karaarslan, and F. Safa Erenay, “Generative AI in Academic Writing: A Comparison of DeepSeek, Qwen, ChatGPT, Gemini, Llama, Mistral, and Gemma,” 2025.

[23] A. Partin et al., “Deep learning methods for drug response prediction in cancer: Predominant and emerging trends,” Front Med (Lausanne), vol. 10, p. 1086097, Feb. 2023, doi: 10.3389/FMED.2023.1086097/XML/NLM.

[24] O. Erdem, K. Hassett, and F. Egriboyun, “Hallucination in AI-generated financial literature reviews: evaluating bibliographic accuracy,” Int J Data Sci Anal, pp. 1–10, Feb. 2025, doi: 10.1007/S41060-025-00731-0/TABLES/2.

[25] K. E. Gumilar et al., “Accuracy and hallucination of DeepSeek and ChatGPT in scientific figure interpretation and reference retrieval,” Jun. 2025, doi: 10.21203/RS.3.RS-6676676/V1.

[26] G. Marvin, N. Hellen, D. Jjingo, and J. Nakatumba-Nabende, “Prompt Engineering in Large Language Models,” pp. 387–402, 2024, doi: 10.1007/978-981-99-7962-2_30/TABLES/2.

[27] W. Zhang and J. Zhang, “Hallucination Mitigation for Retrieval-Augmented Large Language Models: A Review,” Mathematics 2025, Vol. 13, Page 856, vol. 13, no. 5, p. 856, Mar. 2025, doi: 10.3390/MATH13050856.

[28] A. Madaan et al., “SELF-REFINE: Iterative Refinement with Self-Feedback,” 2023, Accessed: Jun.30, 2025. [Online]. Available: https://selfrefine.info/

[29] S. Song, J. Lee, and H. Ko, “Hansel: Output Length Controlling Framework for Large Language Models,” Dec. 2024, Accessed: Jun. 30, 2025. [Online]. Available: http://arxiv.org/abs/2412.14033

[30] R. Jie, X. Meng, L. Shang, X. Jiang, and Q. Liu, “Prompt-Based Length Controlled Generation with Multiple Control Types,” 2024.

[31] B. Butcher, M. O’Keefe, and J. Titchener, “Precise length control for large language models,” Natural Language Processing Journal, vol. 11, p. 100143, Jun. 2025, doi: 10.1016/J.NLP.2025.100143.

